# Development and Implementation of a Core Genome Multilocus Sequence Typing (cgMLST) scheme for *Haemophilus* influenzae

**DOI:** 10.1101/2024.04.15.589521

**Authors:** Made Ananda Krisna, Keith A. Jolley, William Monteith, Alexandra Boubour, Raph L. Hamers, Angela B. Brueggemann, Odile B. Harrison, Martin C. J. Maiden

## Abstract

2.

*Haemophilus influenzae* is part of the human nasopharyngeal microbiota and a pathogen causing invasive disease. The extensive genetic diversity observed in *H. influenzae* necessitates discriminatory analytical approaches to evaluate its population structure. This study developed a core genome MLST (cgMLST) scheme for *H. influenzae* using pangenome analysis tools and validated the cgMLST scheme using datasets consisting of complete reference genomes (N=14) and high-quality draft *H. influenzae* genomes (N=2,297). The draft genome dataset was divided into a development (N=921) and a validation dataset (N=1,376). The development dataset was used to identify potential core genes with the validation dataset used to refine the final core gene list to ensure the reliability of the proposed cgMLST scheme. Functional classifications were made for all resulting core genes. Phylogenetic analyses were performed using both allelic profiles and nucleotide sequence alignments of the core genome to test congruence, as assessed by Spearman’s correlation and Ordinary Least Square linear regression tests. Preliminary analyses using the development dataset identified 1,067 core genes, which were refined to 1,037 with the validation dataset. More than 70% of core genes were predicted to encode proteins essential for metabolism or genetic information processing. Phylogenetic and statistical analyses indicated that the core genome allelic profile accurately represented phylogenetic relatedness among the isolates (*R*^2^ = 0.945). We used this cgMLST scheme to define a high-resolution population structure for *H. influenzae*, which enhances the genomic analysis of this clinically relevant human pathogen.

**Impact statement:** Discriminating *H. influenzae* variants and evaluating population structure has been challenging and largely unstandardised. To address this, we have developed a cgMLST scheme for *H. influenzae.* Since an accurate typing approach relies on precise reflection of the underlying population structure, we explored various methods to define the scheme. The core genes included in this scheme were predicted to encode functions in essential biological pathways, such as metabolism and genetic information processing, and could be reliably assembled from short-read sequence data. Single-linkage clustering, based on core genome allelic profiles, showed high congruence to genealogy reconstructed by Maximum-Likelihood (ML) methods from the core genome nucleotide alignment. The cgMLST scheme v1 enables rapid and accurate depiction of high-resolution *H. influenzae* population structure, and making this scheme accessible via the PubMLST database, ensures that microbiology reference laboratories and public health authorities worldwide can use it for genomic surveillance.

**Data summary:** The *H. influenzae* cgMLST scheme is accessible via https://pubmlst.org/organisms/haemophilus-influenzae. The list of isolate IDs available publicly from pubmlst.org is provided in Supplementary File 1. The pipeline for cgMLST scheme development and validation is published at https://www.protocols.io/private/EF6DB7FE429311EEB8630A58A9FEAC02. All in-house R and Python scripts for data processing and analysis are available from https://gitfront.io/r/user-4399403/ZHt8DArALHcY/cgmlst-hinf/.

## 5. Introduction

*Haemophilus influenzae* is a fastidious Gram-negative commensal coccobacillus which is also an accidental pathogen in humans. It is classified based on the expression of capsular polysaccharides, categorised as six serotypes, a to f, plus the unencapsulated or nontypeable (NTHi). *H. influenzae* exclusively inhabits the human host, primarily the upper respiratory tract (URT), where it resides as a commensal member of the microbiota. However, infections caused by *H. influenzae* can manifest as non-invasive or invasive diseases, including otitis media, sinusitis, meningitis, epiglottitis, orbital cellulitis, and septicaemia [1, 2]. Prior to the implementation of the *H. influenzae* serotype b (Hib) polysaccharide-conjugate vaccine, Hib was the predominant causative agent of *H. influenzae* invasive diseases worldwide. Currently, more than 70% of cases can be attributed to NTHi, which is also the most prevalent *H. influenzae* group [3, 4]. The incidence of these diseases has consistently increased across all age groups in recent years [3, 5, 6]. The circulating invasive NTHi has been reported to be increasingly resistant to multiple antibiotic groups such as beta-lactams, cephalosporins, fluoroquinolones, and macrolides [7–9]. Additionally, vaccines for non-Hib *H. influenzae* are currently unavailable with no vaccine development made in the past 5 years [10–13]. Although surveillance programs exist [14], with most centres in high-income countries incorporating whole-genome sequencing (WGS), there is a lack of standardised, high-resolution methods and nomenclature to describe and delineate important lineages.

Molecular classification of human pathogens is an integral component of microbiological diagnostics and surveillance programmes. Among the most widely employed methodologies is multilocus sequence typing (MLST), which is based on the genetic variability within a set of six to eight housekeeping gene fragments, to categorise bacterial isolates into distinct sequence types (STs). These STs, in turn, can be organised into clonal complexes (CCs) based on similarities in their allelic profiles [15, 16]. Core genome MLST (cgMLST) extends the MLST framework to hundreds of complete core genes (i.e. genes shared among all or nearly all isolates of a species) to assess genomic variation with greater precision [17, 18]. Like conventional MLST, cgMLST treats all allelic differences as a single event, as a mitigation of the effects of recombination on tree branch lengths [19]. Because of this, previous reports have concluded that it is the preferred method for assessing highly recombining organisms, including *H. influenzae* [20, 21]. *H. influenzae*, particularly NTHi, exhibits substantial genomic diversity primarily attributed to horizontal gene transfer (HGT) through transformation and recombination [22]. For example, analyses of 24 *H. influenzae* isolates revealed nearly 48,000 open reading frames which were grouped into 3,100 orthologous gene clusters. Only 1,538 of these clusters were shared among all isolates, forming the core genome [22–24].

Previous investigations into the core genome of *H. influenzae* have yielded insights, with one study culminating in the development of a cgMLST scheme; however, these studies typically included fewer than 500 draft genomes, which has been identified as a threshold for stable paralogous locus detection and a key step in core genome analyses [25–27]. Paralogs are duplicated genes derived from a single gene and their presence can mislead genetic relationships among genomes [28, 29]. Furthermore, neither of the existing cgMLST schemes had been validated or implemented on publicly accessible platforms. Previous research has identified various *in silico* parameters affecting cgMLST precision, the most important of which was the completeness of a cgMLST profile [30]. The BIGSdb software that hosts the PubMLST website periodically scans for new alleles within defined schemes, such as cgMLST. These alleles are stored in the sequence definition (seq-def) database and assigns allele numbers, with the resulting cgMLST profile defined in the same database, ensuring the fulfilment of completeness criteria [31]. At the time of writing, the PubMLST website had implemented cgMLST schemes for various bacterial species, including *Neisseria meningitidis* [32]*, N. gonorrhoeae* [33]*, Campylobacter* sp [34]*, Streptococcus agalactiae*, *S. pneumoniae* [35]*, S. uberis* [36]*, Vibrio cholera* [37]*, V. parahaemolyticus* [38], *Bacillus anthracis* [39]*, B. cereus* [40]*, Burkholderia mallei* [41]*, Acinetobacter baumannii* [42], and *Clostridium perfringens* [43].

This study developed and validated a cgMLST scheme for *H. influenzae*, which was then implemented in the PubMLST database. The scheme is publicly available and can be utilised by public health authorities worldwide to characterise *H. influenzae* genomes in a more standardised manner and with greater detail. It also serves as a tool to improve our understanding of the population biology of the bacterium, particularly the NTHi as this approach is minimally biased by recombination events frequently occurring in this population. In turn, this can be used to aid in vaccine development and contextualise antimicrobial resistance (AMR) spread.

## 6. Methods

### Choosing pangenome analysis tools and dataset compilation

#### Reference genomes were used to develop a computational pipeline combining several pangenome analysis software packages

Several open-source pangenome analysis software packages were available at the time of analyses (May 2023), including Roary [44], PIRATE [45], PanX [46], PGAP [47], PPanGGOLiN [48], MetaPGN [49], PEPPAN [50], chewBBACA [27], and Panaroo [51]. Tools without any bug fixes or updates for 5 years or more were not used for this analysis. Roary and PPanGGOLiN were excluded because, based on a recent report, the accessory genome size was likely to be inflated using these programs [51]. Therefore, four software packages, PIRATE, PEPPAN, chewBBACA, and Panaroo were employed. A *‘*two-step’ approach was implemented to evaluate the optimal pipeline for *H. influenzae* core gene identification:

1. Assessment of the methods and unique features provided by each software (Supplementary Table 1); and,
2. Pangenome analysis of a set of reference genomes using each of the software packages and comparison of the outputs.

The complete workflow and decision-making that followed for each step are published in https://www.protocols.io/private/EF6DB7FE429311EEB8630A58A9FEAC02.

#### Development and validation datasets of high-quality draft genomes

The dataset for this study was compiled from genomes stored in the PubMLST database, accessed on 24^th^ September 2022. All publicly available *H. influenzae* genomes in the database underwent quality checks based on several parameters (Supplementary Table 2) [52]. Genomes not fulfilling these criteria for any of the parameters were individually reviewed for inclusion into the dataset. After the review, 2,397 *H. influenzae* isolates were chosen, and their provenance data were obtained. Their genomes were systematically allocated into two independent datasets, the development (N=986) and the validation dataset (N=1,411). The allocation was conducted based on the distribution of important clinical and epidemiological variables in the dataset, including capsule genotype, isolate source, disease, and region of isolation (Supplementary Figure 1).

### Core gene identification and curation

For the purpose of the scheme generation, core genes were defined as genes present in at least 95% of isolates [17, 32, 53]. After finalising the computational pipeline, PIRATE was mainly used for core gene identification, with the help of Panaroo, because it incorporates additional stages for annotation error correction. Results from the two programs were consolidated by utilising an in-house Python script, available on https://www.protocols.io/private/EF6DB7FE429311EEB8630A58A9FEAC02 and https://gitfront.io/r/user-4399403/ZHt8DArALHcY/cgmlst-hinf/. To increase the sensitivity of paralog detection and their exclusion from the final cgMLST scheme, another pangenome suite, PEPPAN, was employed (Figure 1).

**Figure 1.**
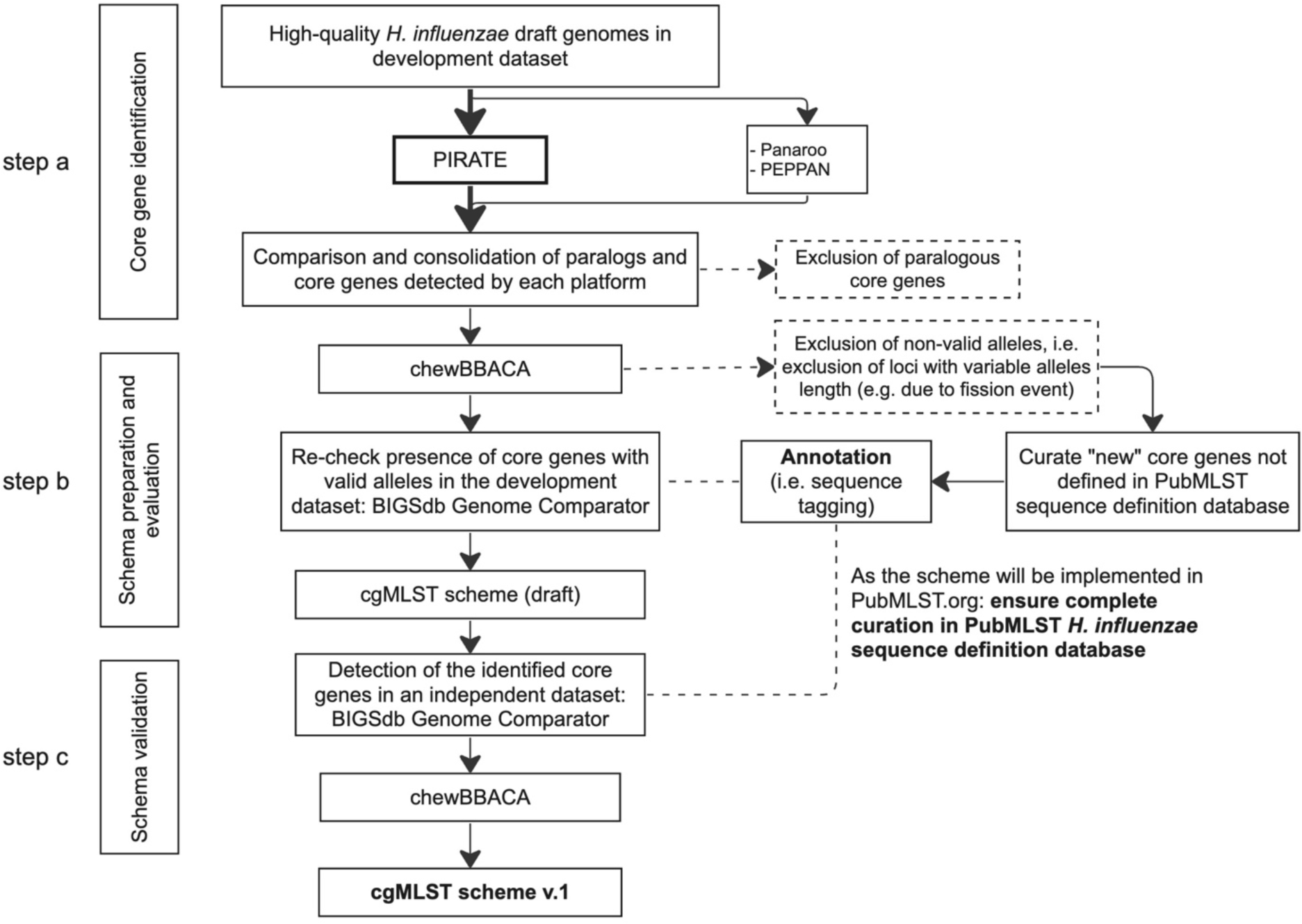
The workflow of cgMLST scheme development and validation.

Each core gene identified in step a (Figure 1) was evaluated for the presence of invalid alleles using chewBBACA with default parameters (Figure 1, step b). Invalid alleles were defined either as: i) sequences with ambiguous characters; ii) a length not divisible by 3; iii) the presence of in-frame stop codon(s); and, iv) sequences that could not be translated due to missing start/stop codons [27]. Valid core genes were subsequently cross-referenced with the PubMLST database to identify genes that had not been previously defined in the database and allow these to be curated in the database. To automatically detect new alleles, the BIGSdb ‘sequence tagging’ function was utilised (90% minimum identity; 70% minimum alignment; and a BLASTN word size of 20) [54]. All defined genes in the PubMLST *H. influenzae* database were assigned a unique locus name starting with “HAEM”, followed by an arbitrary number. Each HAEM locus can be associated with the common gene name, e.g. HAEM1156 corresponds with the *bexA* gene in capsule region I.

### Validation analyses

The initial cgMLST scheme was applied to the validation dataset to confirm that the core genes were still defined as core using a different dataset. This was implemented using the BIGSdb Genome Comparator tool available in PubMLST (default parameters). All genes which remained core after this step would constitute the final *H. influenzae* cgMLST scheme. Additional analyses were performed on these core genes, including allelic variability, functional classification, and intragenic recombination analysis.

To assess the allelic variability of core genes, the total number of alleles and their length were calculated using an in-house Python script (https://gitfront.io/r/user-4399403/ZHt8DArALHcY/cgmlst-hinf/). Functional classification of the core genes was completed using eggnog-mapper (version 2.1.11), using Diamond in blastx mode and HMMER methods, with default parameters [55]. Each gene was assigned a Cluster of Orthologous Genes (COG) category [56]. The COG category was further grouped based on the Kyoto Encyclopedia of Genes and Genomes (KEGG) BRITE functional hierarchy system [57]. Lastly, intragenic recombination analysis was conducted using the Pairwise Homoplasy Index (PHI) method with PhiPack software. PHI is a measure of recombination by assessing the incompatibility between each site and its downstream sites within a global alignment. To evaluate its statistical significance, it calculates a p-value by comparing the observed PHI statistic to a normal distribution derived from the expected mean and variance [58, 59].

### Phylogenetic analysis

A maximum-likelihood (ML) tree was generated based on the core genome nucleotide alignment of 1,376 genomes in the validation dataset using RaXML (version 8) and ClonalFrameML, which accounts for recombination events [60, 61]. The tree was then annotated with the set of metadata detailed below, along with the method(s) employed for their retrieval:

1. **Core genome cluster (CGC) at 500, 200, and 50 allelic mismatches**. Every genome in the dataset received a core genome sequence type (cgST) based on the allelic profile of the 1,037 core genes contained within the scheme. For a cgST to be assigned, a maximum of 25 genes can be missing. This threshold was set because the scheme was applied in draft genomes which may lack complete annotation.

All genomes with a cgST assigned were further grouped into CGCs using a single-linkage clustering method with the pre-determined similarity thresholds at 500, 200, and 50 allelic mismatches. Each member of the CGC group has a maximum of the specified allelic differences with at least one other member of the group.

1. **Capsule type based on genome sequences** (i.e. capsule genotype) as assigned by the Hicap suite software [62].
2. **Clonal complexes (CC)** were defined using the globally-optimised eBURST (goeBURST) algorithm implemented in PHYLOViZ 2.0.
3. **Pathotype clade classification system for NTHi**. Previous studies on NTHi population structure revealed six distinct clades (Clade I-VI), based on the presence or absence of 17 accessory genes (Supplementary Table 3), which was reported to reflect the phylogeny based on the core genome single nucleotide polymorphisms (SNPs) [26, 63, 64]. The presence of these genes was concluded by conducting BLAST search to genomes in the dataset, with the following parameters: 90% identity, 70% alignment, and BLASTN word 20.
4. **Biotype** based on the presence or absence of genes (Supplementary Table 4) encoding three metabolic enzymes: ornithine decarboxylase (ODC), urease, and tryptophanase [65, 66].

Additionally, a minimum-spanning tree (MST) based on the core genome allelic profile was constructed, utilising the GrapeTree plugin on PubMLST isolate database [54, 67].

### Relationships between the cgMLST scheme pairwise allelic mismatch and ML tree branch length

Genetic relatedness among *H. influenzae* isolates was assessed based on their core genome by two approaches either using the core genome allelic profile or the nucleotide sequence identity. To evaluate congruence between these approaches, scatter plots were generated, incorporating a correlation test and simple linear regression analysis with Ordinary Least Square (OLS) method. Distance matrices based on cgMLST scheme pairwise allelic mismatch and core genome alignment ML tree branch length were produced using the Genome Comparator plug-in on the PubMLST website and an in-house Python script (https://gitfront.io/r/user-4399403/ZHt8DArALHcY/cgmlst-hinf/). Matrices were converted into a frequency table and a scatter plot was generated. If at least one group showed non-Gaussian distribution, the Pearson correlation test was employed for calculating the p-value. Lastly, regression analysis was conducted to obtain a mathematical function explaining the relationship between the two methods, as well as the coefficient of determination (R^2^) which measures the goodness-of-fit of the function. These steps were conducted with R Statistical Software and Python statistics packages. All scripts are available on https://gitfront.io/r/user-4399403/ZHt8DArALHcY/cgmlst-hinf/.

## 7. Results

### *H. influenzae* genomes from PubMLST database used for cgMLST scheme development and validation

The reference genome dataset was comprised of 14 complete reference *H. influenzae* genomes, 11 of which were listed in KEGG Organisms: Complete Genomes database (https://www.genome.jp/kegg/catalog/org_list.html accessed 26 September 2022). Three additional clinically important complete genomes were included after a publication search through NCBI PubMed [68, 69]. All associated provenance data and complete genome sequences are available from PubMLST (Supplementary Table 5).

There were 986 draft genomes in the development dataset that fulfilled the quality check criteria; however, this number was reduced to 921 genomes after initial pangenome analysis with PIRATE. The 65 excluded genomes exhibited high duplication events and corresponded to the outliers in the box plot for genome length (Supplementary Figure 2) of the original development dataset, and likely represent sequencing errors or lower assembly quality. Therefore, the pangenome analysis was rerun using 921 draft genomes in the development dataset, which identified 1,063 core genes for the first draft of the cgMLST scheme. Subsequently, genomes in the validation dataset (N = 1,411) were annotated for these core genes and genomes with < 95% of the core genes annotated were excluded (N = 35), resulting in the final validation dataset of 1,376 genomes. There was no notable difference in the distribution of other variables between the initial and final datasets (Supplementary File 1).

*H. influenzae* isolates in both datasets were globally distributed; however, most isolates originated from North America (356/921 and 548/1376) and Europe (431/921 and 547/1376). The two datasets also showed a similar temporal distribution, with most isolated after 2016. There were more clinical isolates from invasive disease cases (i.e. bacteraemia, meningitis, and other invasive) in the development dataset, which was reflected in a similar proportion of non-typeable (542/921) and typeable (379/986) in the dataset. This was done to ensure that the developed schema would be applicable to all typeable *H. influenzae* as well as the NTHi (Figure 2 (i) – (iv)). Based on several metrics, genomes in both datasets were of good quality, with the majority (893/921 and 1346/1376) having less than 150 contigs and an N50 parameter of more than 23,000.

**Figure 2.**
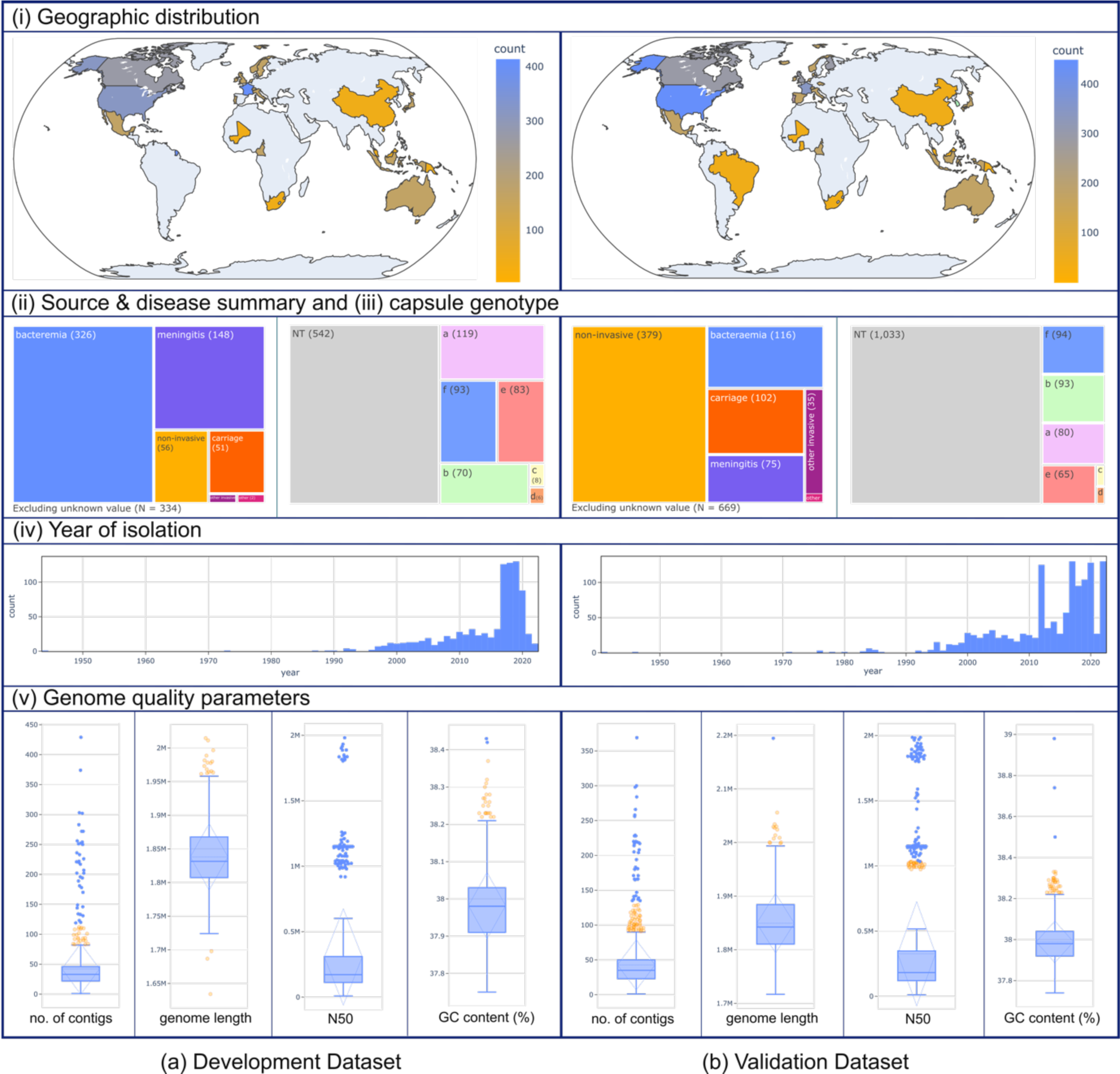
Characteristics of the datasets and genomes employed for developing the *Haemophilus influenzae* cgMLST scheme, (a) development (N = 921) and (b) validation (N = 1,376).

### 1,037 core genes in the validated cgMLST scheme are implicated in important cellular pathways

A total of 1,248 non-paralogous core genes were identified in the combined pangenome analysis with the three tools, PIRATE, Panaroo, and PEPPAN, with 144 paralogous core genes excluded at this stage (Figure 1, step a). Only 18 of these genes were not previously defined in the PubMLST *H. influenzae* sequence definition database and were therefore added. After filtering for invalid alleles (Figure 1, step b), a total of 185 genes were no longer defined as core, resulting in 1,063 genes included in the cgMLST scheme draft (Supplementary File 2). Lastly, 1,037 genes remained core after validation with the validation dataset (Figure 1, step c), excluding only 26 genes, 25 of which were present in 90 – 95% of the genomes (Supplementary File 2).

A total of 1,024 core genes in the scheme were assigned an NCBI COG category, 37 of which had more than one COG category defined [56]. Among the core genes with one COG, 149 (14%) had “Function Unknown”, or denoted as “S”, as the COG category. After re-assessment, 95 had at least one specific, non-S COG based on the orthologous (OG) hit from any higher taxonomy levels. The remaining genes were curated manually using the KEGG orthology classification system or protein product-based search on the Protein Families Database (v 95.0) or both (Supplementary File 3). For the 13 genes without initial COGs category, re-searching using the HMMER method resulted in identical results. These were categorised as having an unknown function and assigned the “S” COG category (Figure 3a).

**Figure 3.**
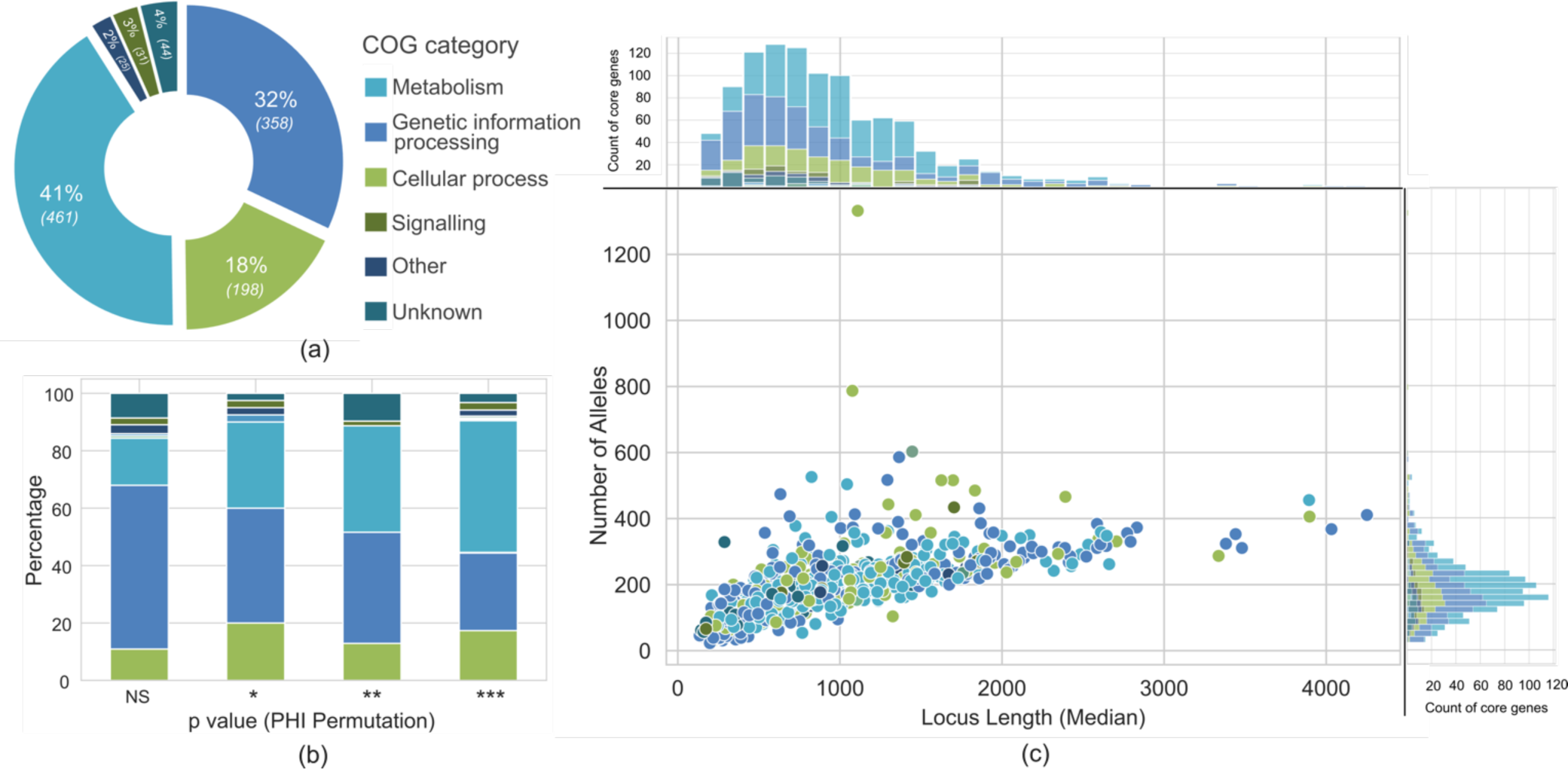
Functional classification, recombination, and allele variability analysis of *Haemophilus influenzae* core genes in the cgMLST scheme. (a) Functional classification was achieved with eggnog-mapper, assigning the Cluster of orthologous genes (COG) category for each core gene. (b) Intragenic recombination analysis of each core gene based on pairwise homoplasy index (PHI) permutation p-value. Core genes were grouped based on this p-value: NS non-significant; * .01 < p-value < .05; ** .001 < p-value < .01; *** p-value < .001. The proportion of each COG category within each p-value group was calculated and shown as the coloured stacks in the bar graph. (c) The number of alleles and their length variation were counted for each core gene. The median locus length and the allele count were plotted and coloured based on the COG category. The upper panel showed the distribution of median locus length and on the right panel, the distribution of allele counts; both were coloured in accordance with the COG category.

Nearly 90% (909/1,037) of the genes within the cgMLST scheme exhibited evidence of intragenic recombination events, as indicated by a p-value of < 0.05 in PHI statistics. When assessing the percentage of recombination events in each COG category, the majority (73/128) of core genes with non-significant PHI statistics belonged to the Genetic Information Processing group (Figure 3b). Conversely, nearly all core genes in the Metabolism group (405/461) displayed a p-value of < 0.05.

The number of alleles and median length of each core gene were correlated (Figure 3c). HAEM0191 (hypothetical protein) and HAEM1295 (outer membrane protein P5) exhibited a higher number of alleles compared to other genes of similar median locus length (1332 and 787 alleles, respectively).

### Clustering groups of *H. influenzae* genomes based on pairwise allelic mismatches of the core genes reflected their phylogenetic relationship

Within the validation dataset encompassing 1,376 genomes, 1,320 (95.9%) had a cgST assigned, and thus could be classified into CGC groups at different similarity thresholds (Supplementary File 1). The rest of the genomes without cgST still had an assigned allelic profile for a substantial number of their core genes, ranging from 989 to 1,011 (95.4 – 97.5%), out of the 1,037 core genes in the scheme.

The population structure of *H. influenzae* can be systematically explored using hierarchical core genome clustering at different thresholds. This hierarchical classification, derived from the allelic profiles of the core genome, effectively mirrors the phylogenetic relationships among the isolates (Figure 4a).

**Figure 4.**
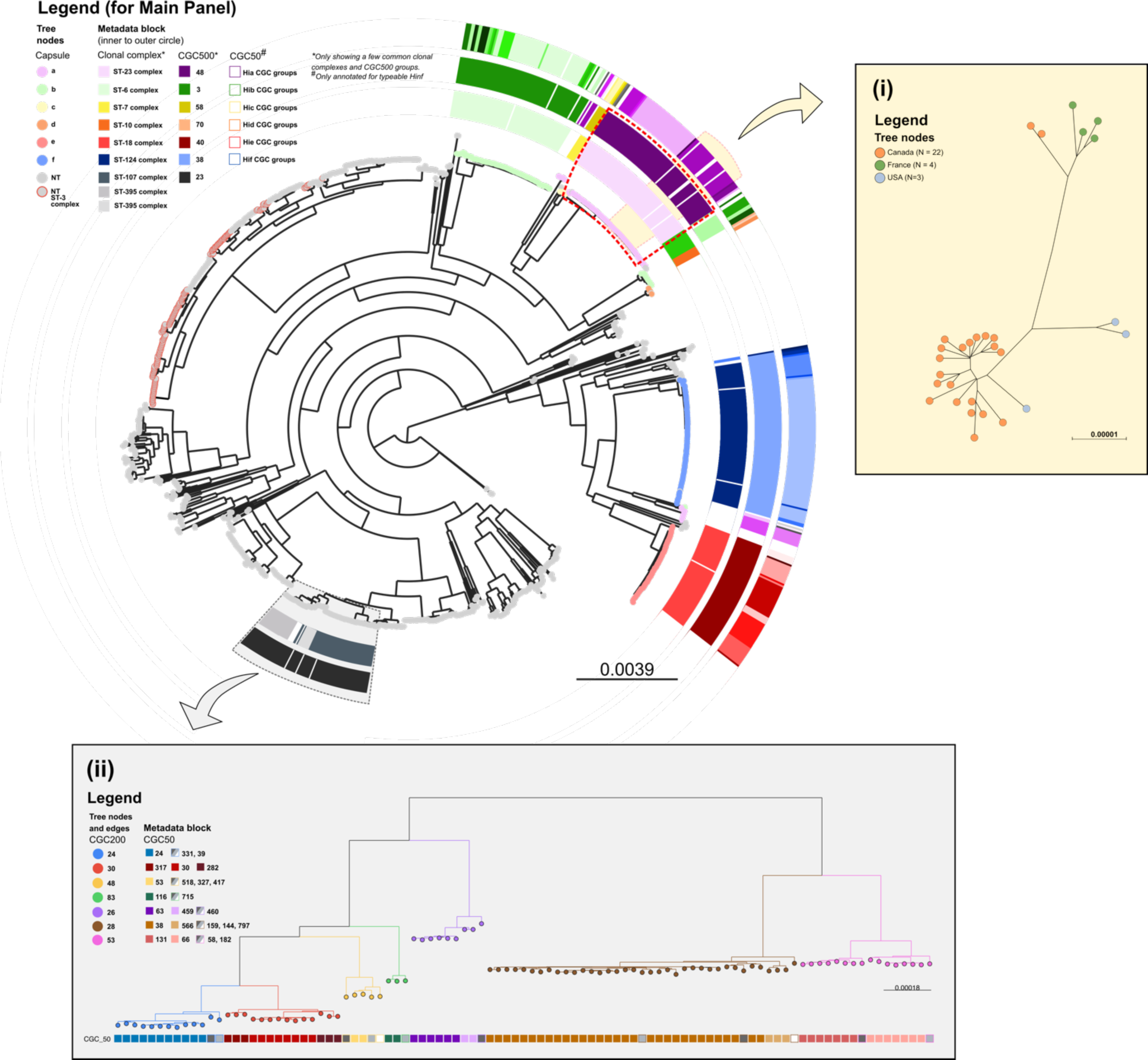
Population structure of 1,376 *H. influenzae* genomes from the validation dataset. **Main panel:** a maximum-likelihood phylogenetic tree generated from a concatenated core gene nucleotide sequence alignment. Tree nodes were coloured by capsule type. Each capsule type clustered together, except for *H. influenzae* type b (Hib). The innermost metadata block was a clonal complex (CC) assignment, which corresponded well with the middle metadata block, representative of the core genome cluster group at the 500 allelic mismatches threshold (CGC500). This correlation was evident for capsulated/typeable *H. influenzae*, but not for NTHi. The outermost metadata block was the core genome cluster group at 50 allelic mismatches threshold (CGC50), which allowed a more granular distinction of *H. influenzae* clusters. The hierarchal clustering at multiple thresholds was able to reflect the structure of the phylogeny. **Subpanel (i):** A subset of the ML tree in yellow highlight, consisting of 29 *H. influenzae* type a (Hia) isolates CGC50 group 15. Isolates in this group were predominantly from North America, with 20 from Canada clustering closely, a pattern reported previously by Topaz et al [70]. **Subpanel (ii):** The hierarchical structure of the ML tree was also represented using CGC based on multiple thresholds for NTHi. For example, a subset of the ML tree highlighted in grey comprises 97 NTHi within the CGC500 23. Tree edges and nodes were coloured based on the CGC200 groups, which are congruent with the tree topology. The metadata block shows CGC50 groups within each CGC200 group. The CGC50 groups with greyscale colour was a singleton within that corresponding CGC200 group.

Typeable *H. influenzae* isolates sharing the same capsule genotype exhibited clustering in both phylogenies, consistent with their close genetic relatedness. Several CGC groups, spanning various allelic mismatch thresholds, aligned with specific capsule genotypes (Figure 4a). For instance, among *H. influenzae* type a (Hia) isolates, four distinct lineages were delineated, each with its own CGC group at 500 allelic mismatches (CGC500) – namely, CGC500 48, 46, 79, and 81 – which correlated with the CC assignments (Supplementary Table 6). The largest Hia lineage, CGC500 48, which corresponds to the ST-23 complex, harboured a significant subgroup, mainly comprised of isolates originating from Canada. This subgroup was clustered together at 50 allelic mismatch thresholds (Figure 4 yellow highlight and subpanel i). A similar pattern emerged within the Hib in which there were three distinct lineages in the ML tree, two of which correlated with the CGC group at 500 allelic mismatches and their respective CCs (Supplementary Table 6). The smallest lineage consisted of a single isolate (isolate ID 5105) of ST 464 and this isolate did not have CC or CGC groups assigned. At the time of writing (October 2023), there is only 1 other genome in the PubMLST database with this ST (isolate ID 15982).

The NTHi isolates were much more diverse, with 110 CGC 500 groups, only 8 of which contained at least 30 isolates in the group (Supplementary Figure 3). Nevertheless, a correlation between CGC grouping and the *H. influenzae* phylogeny was also evident among NTHi. To exemplify this, a subset of the ML tree incorporating 97 NTHi within the CGC500 23 group was highlighted (Figure 4 grey highlight and subpanel (ii)). Within this group, there were 7 clusters at 200 allelic mismatches threshold (CGC200), and this clustering was highly congruent with the tree topology. However, while typeable *H. influenzae* isolates sharing the same CC exhibited clustering congruence in the phylogeny, this was not observed in the case of NTHi. For example, the ST-3 complex exhibited a scattered distribution in the tree (Figure 4 grey nodes with red line). This pattern was observed in 19 out of 57 CCs within the NTHi population in the validation dataset (Supplementary Table 6). Conversely, the NTHi CC assignments did not consistently align with the CGC500 grouping. For instance, within the CGC500 23 group mentioned earlier, three different CCs were identified (Supplementary Table 6). Therefore, the CGC groups offered a more precise representation of the NTHi phylogenetic structure.

Although, in general, there were specific CCs for each *H. influenzae* serotype, this was not always the case. Within the ST-124, ST-210, and ST-422 complexes, there were NTHi and *H. influenzae* type f (Hif) isolates. Conversely, the CGC500 group assignment for these isolates was consistent with both the serotype and phylogeny, also showing the expected advantage of cgMLST compared to 7-loci MLST. Taking this into account, a few NTHi isolates (N = 5) were observed within the encapsulated clusters, with the CGC500 group assignment identical to the encapsulated isolates (Supplementary File 1 and Supplementary Table 6). This marked the possible rare events of capsule loss of previously encapsulated *H. influenzae*.

The ability of the core genome allelic profile within the scheme to depict the phylogenetic relationships in *H. influenzae* was quantitatively assessed (Figure 5). This assessment was done by comparing the number of allelic mismatches with the branch length value from the ML tree for each pair of genomes in the validation dataset. The coefficient of determination (R^2^) of 0.945 indicates that 0.945 of variance in branch lengths, in the context of an ML tree unaffected by recombination events, can be elucidated through the allelic pairwise mismatches observed across the 1,037 core genes. A Spearman correlation test also yielded a statistically significant p value of < 0.001.

**Figure 5.**
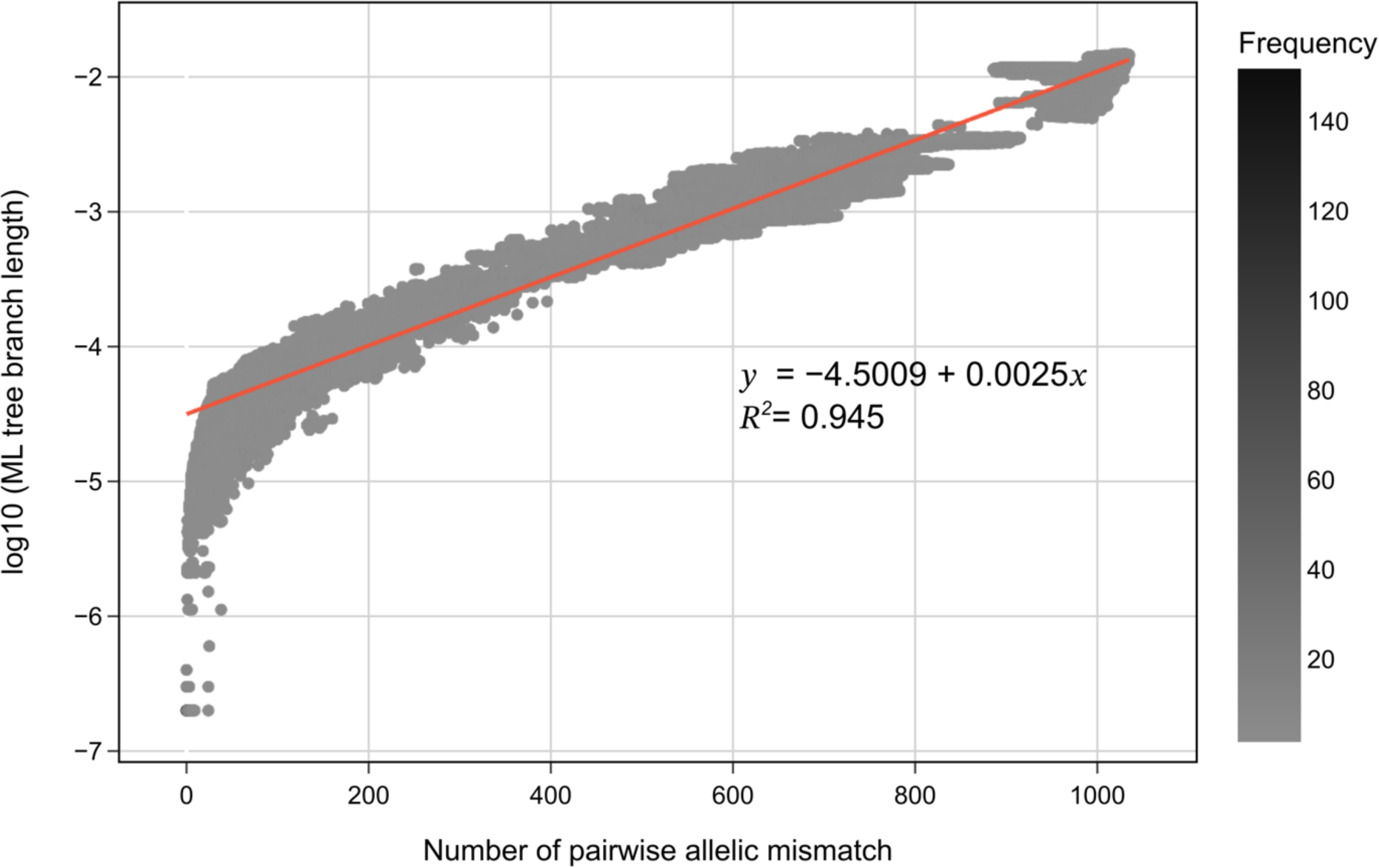
Comparing pairwise allelic mismatch of core genes in the cgMLST scheme with the log10 branch length values from the ML tree, implemented in the validation dataset (N = 1,376). The branch length value was measured for each unique pair of genomes in the dataset. The more closely related genomes in the pair, the lower the allelic mismatch and log10 branch length value; and *vice versa*. The log10 branch length as a function of pairwise allelic mismatch was calculated using an Ordinary Least Square (OLS) method and the adjusted R^2^, a coefficient of determination of the defined function, was also measured.

Lastly, we assessed two previously utilised classification systems aimed at characterising the population structure of *H. influenzae*, to determine the extent of their concordance with both the cgMLST scheme and the overall phylogeny of this bacterial species. First, the pathotype clade classification system, specifically applicable to NTHi, was concluded and annotated in the ML tree of genomes in the validation dataset (Supplementary Figure 3). Out of 1008 NTHi isolates, 160 could not be assigned to any of the six specified clades. Additionally, although the clade classification system followed the phylogeny topology to some extent, within each identified clade, the genetic relatedness among genomes varied greatly. For example, the median pairwise allelic mismatch in clade VI was 991 (minimum-maximum = 0-1028) and in clade II the median was 640 (minimum-maximum = 0-766) (Supplementary Figure 4). Second, we examined the biotype classification system, which relied on the distinct production of three specific enzymes, each directly corresponding to the presence of a particular gene. None of the eight biotypes demonstrated a clear correspondence with specific CGC groups at any threshold, the underlying phylogenetic structure, or any discernible clinical or demographic characteristics (Supplementary Figure 5).

## 8. Discussion

At the time of writing, there were over 560,000 prokaryote genome assemblies in the NCBI Genome Library https://www.ncbi.nlm.nih.gov/genome/microbes/). The accessibility and relative affordability of high-throughput sequencing technologies have resulted in an unprecedented amount of available bacterial genome data [71]. cgMLST is a gene-by-gene typing approach able to assess genomic variation within a bacterial species or genus by utilising whole-genome sequencing data. The gene-by-gene approach has several advantages compared to other methods for population genetic evaluation. First, it does not require a reference genome and second, it focuses on functional protein-coding genes. More importantly, this approach treats any allelic changes as single events, a feature useful for highly recombining organisms such as *H. influenzae.* With 1,037 core genes included in the cgMLST scheme, the scheme can be used to generate a high-resolution population structure largely independent of recombination artefacts.

There are variations in the reported number of *H. influenzae* core genes in published literature. Hoggs *et al.* and Eutsey *et al*. found a range of 1,450 to 1,485 genes shared by 100% of genomes, commonly referred to as “hard core” genes [24, 72]. In 2019, Pinto, *et al.* examined the pan-genome of over 200 NTHi genomes, identifying 1,400 genes shared by at least 95% of the genomes [26]. Rajendra KC *et al*. conducted a similar pangenome analysis in 568 NTHi genomes but found only 853 core genes from the total of 12,249 pan-genes [73]. Gonzalez-Diaz and colleagues revealed a spectrum of 1,470 to 1,627 core genes for each *H. influenzae* capsule type, with a total of 10 to 234 genomes in each capsule group. However, when combined (N = 800 genomes) the core genes shared by all capsulated *H. influenzae* was 1,037 [74]. Several factors are known to cause variations in pangenome analyses. First, core genome size estimation decreases as more genomes are integrated into the analysis until it eventually reaches an asymptote [72]. Second, different approaches and thresholds, used for clustering sequences into the same orthologous group, can either over- or under-estimate core genome size. In turn, these are influenced by the quality of genome sequencing and assembly [50, 51]. Lastly, core genes, estimated by a pangenome analysis, do not automatically exclude putative paralogous genes [27–29]. In developing the cgMLST scheme, we incorporated steps to control for these factors by excluding low-quality genomes based on a set of predetermined criteria and utilising different pangenome analysis tools, particularly to increase the sensitivity of detecting potential paralogous genes. These are all in line with our main aim for the scheme to be used as a robust typing method that reflects the genealogical relationship of *H. influenzae*.

The *H. influenzae* genome clustering based on cgST showed congruence with the genealogy reconstructed using the gold standard ML methods from the core genome nucleotide alignment. Based on the clustering at 500 allelic mismatches, we identified four Hia lineages as described by Topaz et al [70]. They also reported a sublineage mostly isolated from North America, a finding replicated in our study when using the cgST clustering at 50 allelic mismatches [70]. Another report from the U.S. surveillance program discovered two main Hib lineages, also delineated in two different CGC500 groups [75]. However, we identified one Hib of ST 464 (PubMLST id: 5105) located away from the two lineages, indicating a third Hib cluster. In addition, two major Hif lineages clustered at 500 allelic mismatches. A study by Gonzalez-Diaz *et al.* reported only one of these lineages and found three arbitrary clades, which were not discovered in our study (REF). This difference likely stemmed from variations in approaches used to describe Hif population structure (all *H. influenzae* vs. Hif-specific core genome) and in the methodology for generating the phylogeny i.e. core genome nucleotide alignment vs. core SNP using a reference genome [74]. Generally, the cgST clustering at 500 allelic mismatches corresponded to CC assignment, with some exceptions. Some discrepancy was expected as there are 150 times more genes in the cgMLST scheme, compared to MLST, which only has 7, resulting in higher resolution for depicting the population structure. However, the overall correlation between CC and CGC500 group showed that 7-gene MLST remains useful for describing a broader level of genetic relatedness of *H. influenzae* isolates when WGS is not possible.

The five NTHi isolates located within the encapsulated clusters in the ML tree, with their CGC500 group assignment identical to the encapsulated isolates, may signify rare capsule loss events. In 2019, the first case report of a Hib isolate from an invasive case exhibiting loss of capsule expression was reported [76]. Although Potts *et al.* observed a similar finding in the same year through population genetic study of the US *H. influenzae* collection from surveillance programs [75], the implementation of cgMLST and the subsequent cgST clustering will circumvent the need to construct phylogeny in order to observe such phenomena.

Finally, our cgMLST scheme was compared with two existing classification systems for *H. influenzae.* The first and oldest, ‘biotyping’, was utilised prior to the development of molecular techniques in microbiology and relied on the differential production of tryptophanase (indole test), urease, and OCD [77]. Slotved *et al.* demonstrated that the production of these enzymes can be inferred from the molecular detection of the genes encoding them, although their presence/absence did not reflect the phylogeny, which we also demonstrated here [66]. The second was proposed more recently and focused on the NTHi, the NTHi clade typing. This typing system was developed based on core genome SNPs, which was defined as “the portion of the reference genome (isolate 86-026NP, PubMLST id: 5068, NCBI RefSeq GCF_000012185.1) that could be aligned against all of the other sequences” [64, 73]. In two previous studies, each NTHi isolates were assigned to one clade with isolates in clade I, IV, and V forming a monophyletic group in the phylogeny [63, 64]. These findings were not replicated in the current study. Our results suggested that biotyping and NTHi clade typing had a limited discriminatory power and precision for describing *H. influenzae* phylogenetic relatedness [65]. The cgMLST method utilises more genes and a gene-by-gene approach, and can depict a high-resolution population structure free from reference bias.

In conclusion, the cgMLST scheme proposed here provides a high-resolution representation of *H. influenzae* population structure and is a reliable and accessible method to use for characterising phylogenetic relatedness between *H. influenzae* isolates that can be utilised by microbiology reference laboratories and public health authorities worldwide. This represents a significant breakthrough in typing methodologies, particularly for the NTHi population as the predominant cause of *H. influenzae* invasive disease. Moreover, the thorough characterisation of the bacterium core genome serves as a valuable resource for enhancing molecular diagnostics and advancing vaccine development.

## 9. Author statements

### 9.1 Author contributions

M.C.J.M. and O.B.H. designed the study and wrote the manuscript with M.A.K.; M.C.J.M., O.B.H., A.B.B., and R.H. supervised the activities; M.A.K. and A.B.B. curated the draft genomes; A.B.B. and A.B. identified central genotypes for *H. influenzae* clonal complexes description; M.A.K. performed the pangenome analyses, created all figures, curated loci in cgMLST schemes; M.A.K and W.M. wrote in-house R and python scripts; K.A.J. implemented the cgMLST scheme in the PubMLST database.

### 9.4 Ethical approval

Not applicable

### 9.5 Consent for publication

Not applicable

## 9.6 Acknowledgements

The authors wish to thank Prof. Susanna Dunachie and Prof. Samuel K. Sheppard for their valuable advice on the work.

