## Supplementary Figures for "Development and Implementation of a Core Genome Multilocus Sequence Typing (cgMLST) scheme for *Haemophilus* influenzae"

**Supplementary Figure 1**

First (Step 1), the genomes were classified based on the combination of “source” and “disease” provenance data, resulting in seven groups: blood/bacteremia, CSF/meningitis, blood/other invasive diseases, any source/pneumonia, any source/carriage, any source/non-invasive disease, and missing value. Excluding the ones with the missing value, half of the genomes for each group were chosen using a Python randomiser module, resulting in 536 genomes included in the development dataset. These genomes were then evaluated for their capsule type distribution, showing a skewed type b and NTHi distribution. Therefore, the second step (Step 2) aimed to add more non-type b capsulated *H. influenzae*. Excluding the NTHi and type b *H. influenzae* genomes and all isolates included in the first step, a subset of genomes from each capsule group was chosen. This subset is proportionate to the original distribution of each capsule type in the overall dataset. At this stage, a total of 733 genomes were included and their geographical distribution showed a skewed representation from Europe. Consequently, the last step (Step 3) intended to add more genomes from continents other than Europe, with a proportion as close as the overall dataset. Following this process, there were a total of 986 *H. influenzae* genomes for the development dataset.

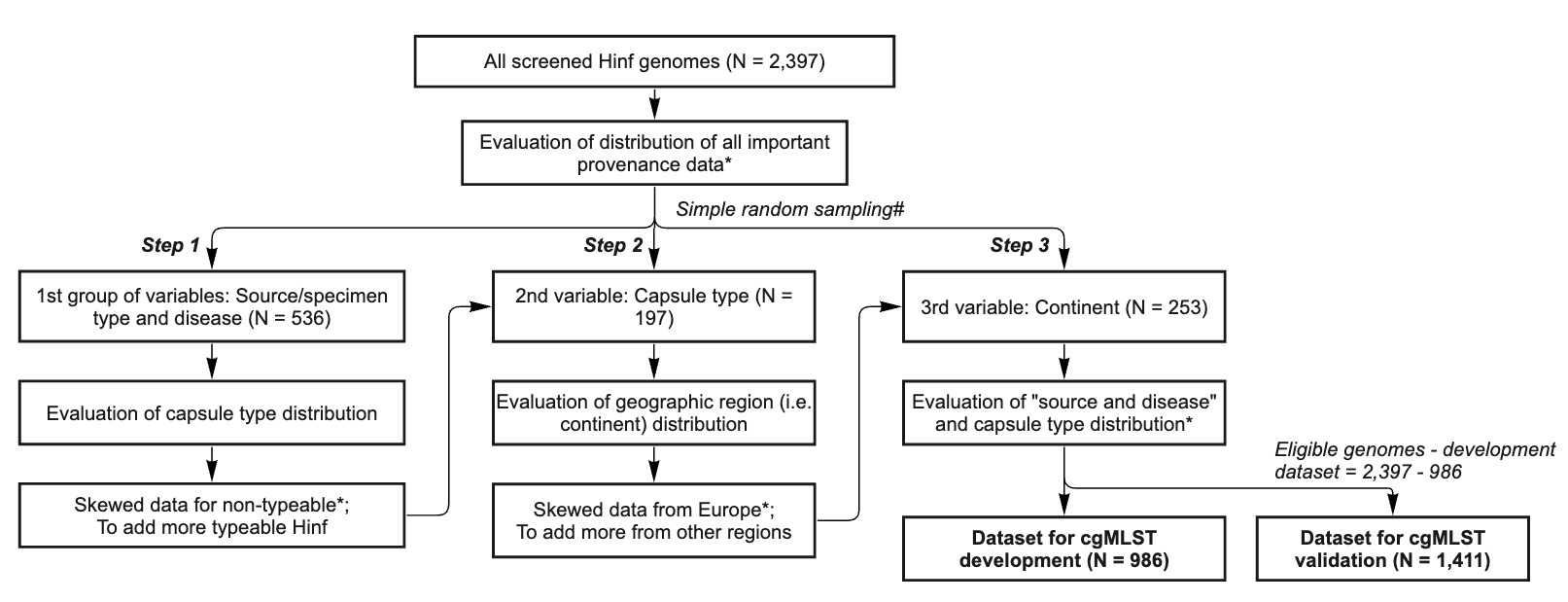

*Importance provenance data are source/specimen type, disease, serotype based on genotype (i.e. capsule type), and geographic location (i.e. continent).

#Steps for selection were done consecutively. For each step, samples were selected through a simple random sampling method from the pool of 2,397 Hinf genomes while considering that any selected sample from the previous step would not be included.

**Supplementary Figure 2**

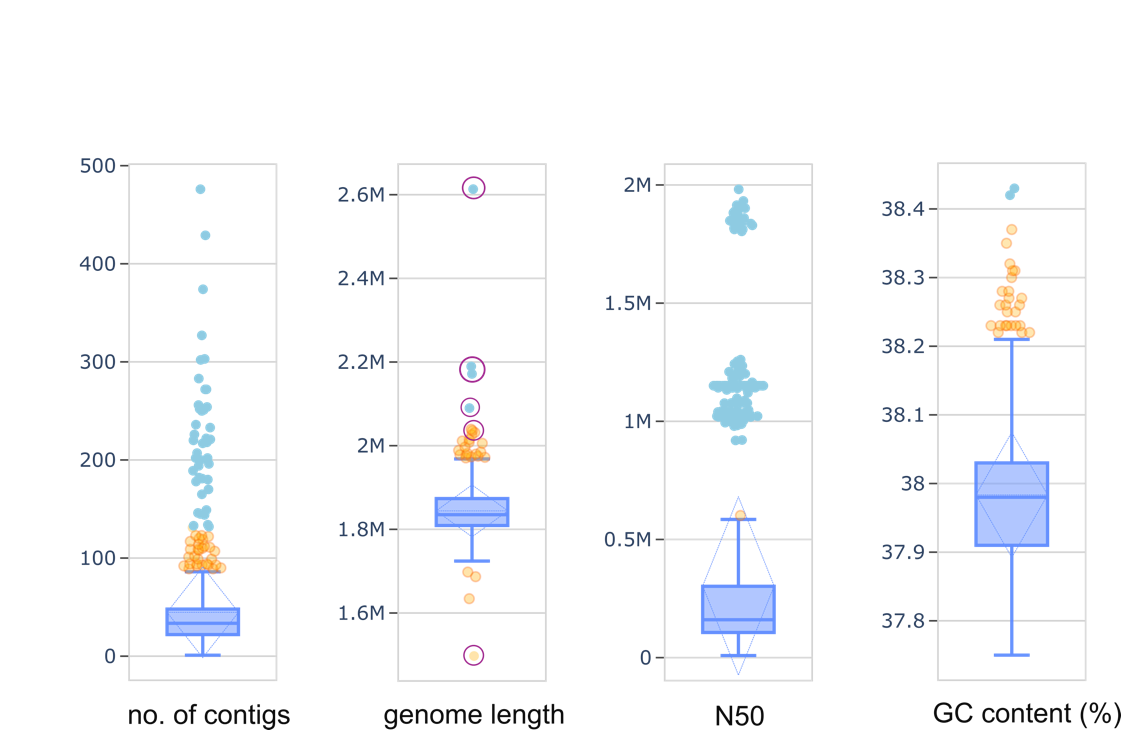
Genome quality metrics of the original development dataset consisting of 986 *H. influenzae* draft genomes. Genomes excluded from this dataset corresponded to the outliers in the “genome length” box plot.

**Supplementary Figure 3.1**

The maximum-likelihood tree from core genome alignment of 1,376 *H. influenzae* genomes in the validation dataset. Tree nodes with black colour were encapsulated isolates. For NTHi, tree nodes were coloured with the CGC500 group, if the group consisted of at least 30 isolates; otherwise, the nodes were coloured as grey. Metadata block shows different NTHi pathotype clades.

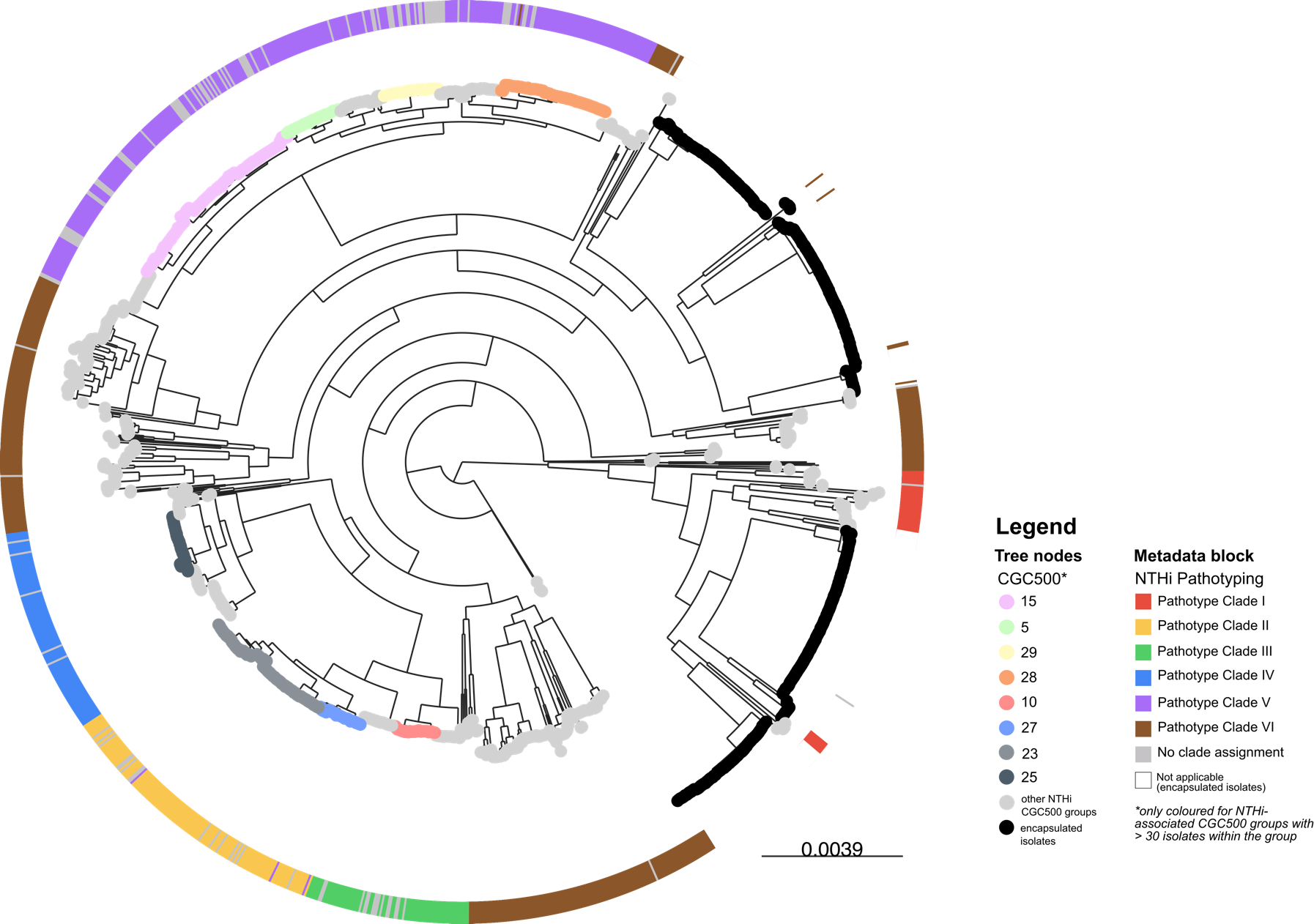

**Supplementary Figure 3.2**

A minimum-spanning tree based on core genome profile, showing different CGC500 groups clustered together.

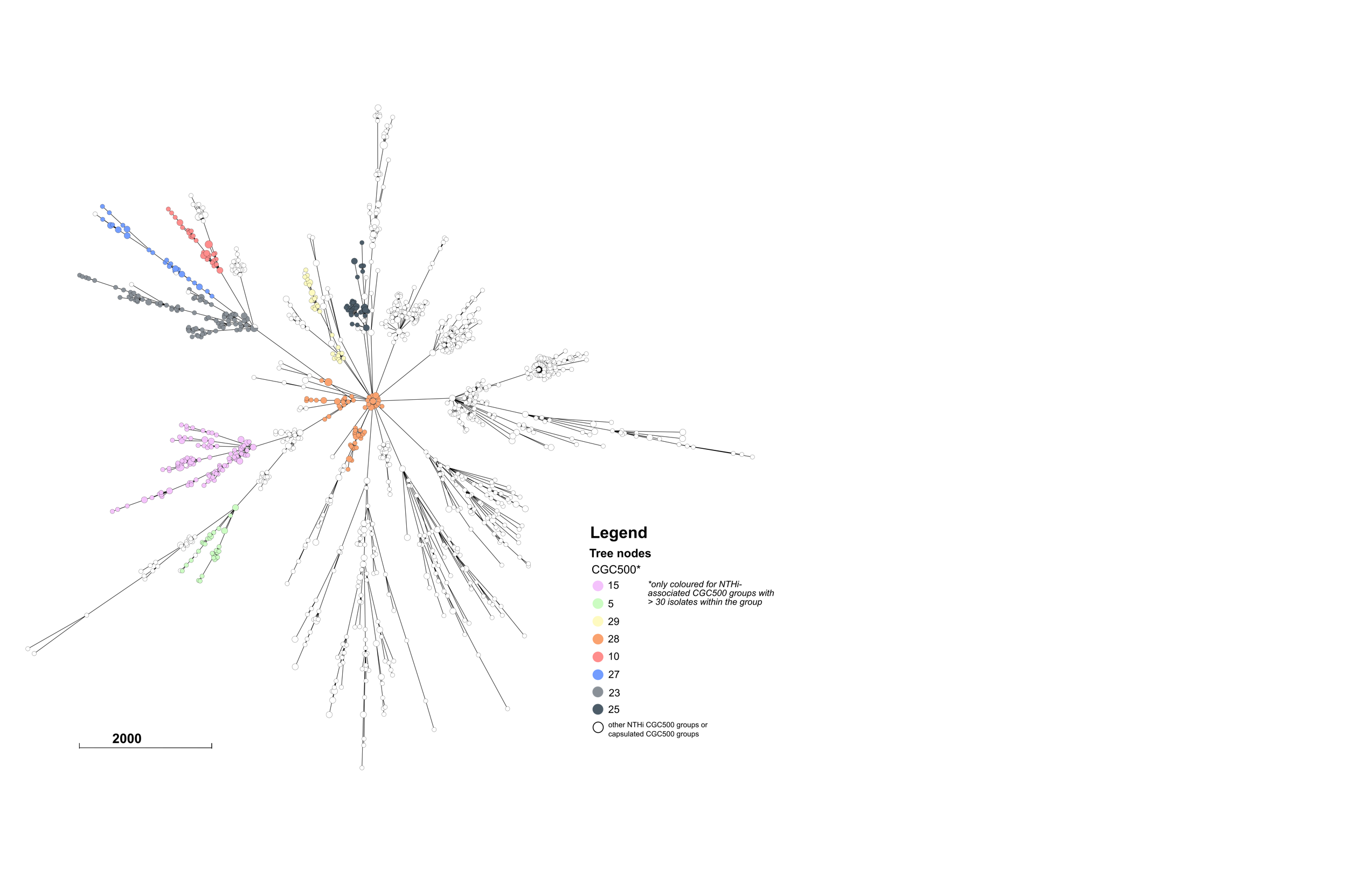

**Supplementary Figure 4**

Variation of genetic relatedness among NTHi isolates within the same pathotype clade. Genetic relatedness was reflected by the number of allelic mismatches for each possible combination of paired isolates (i.e. pairwise allelic mismatch).

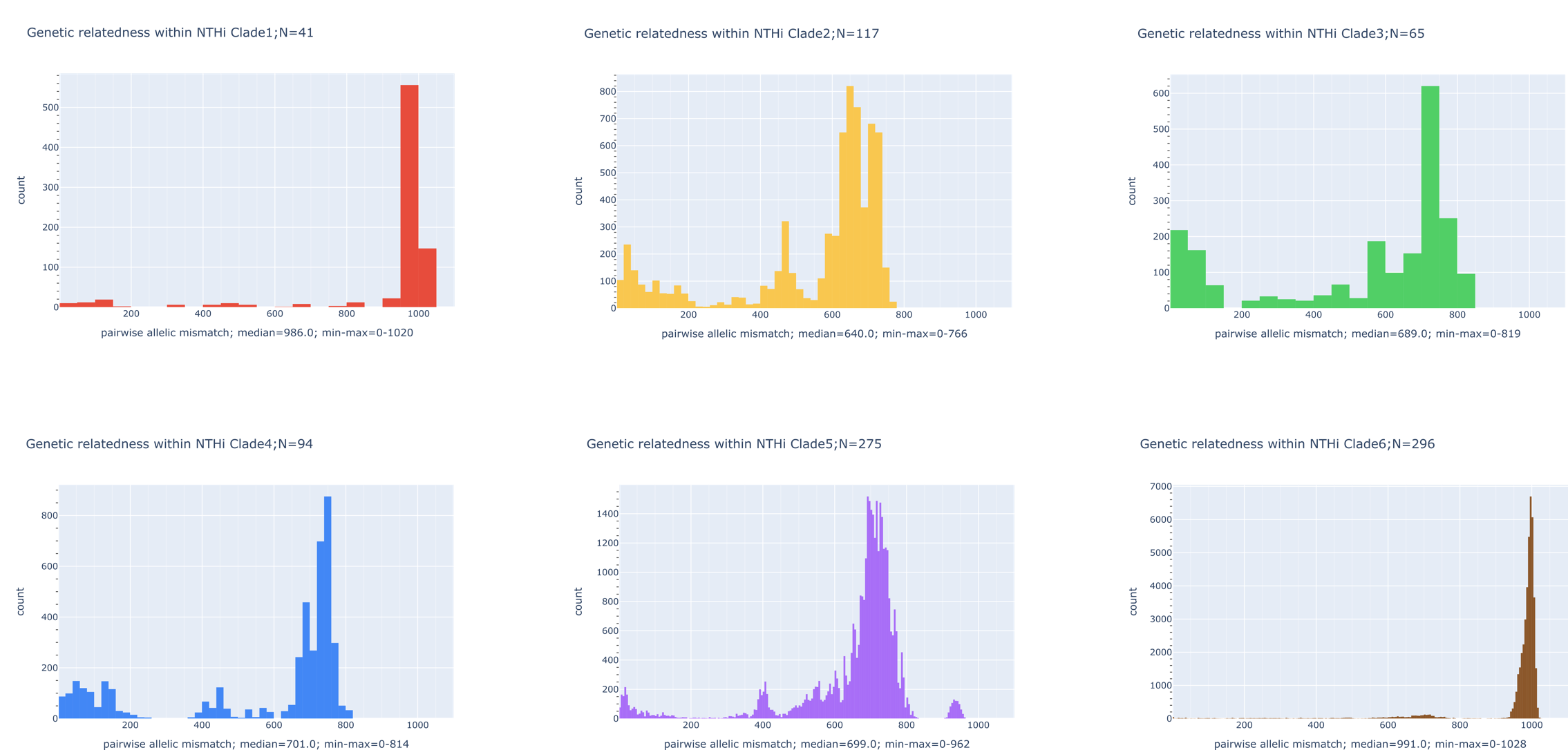

**Supplementary Figure 5**

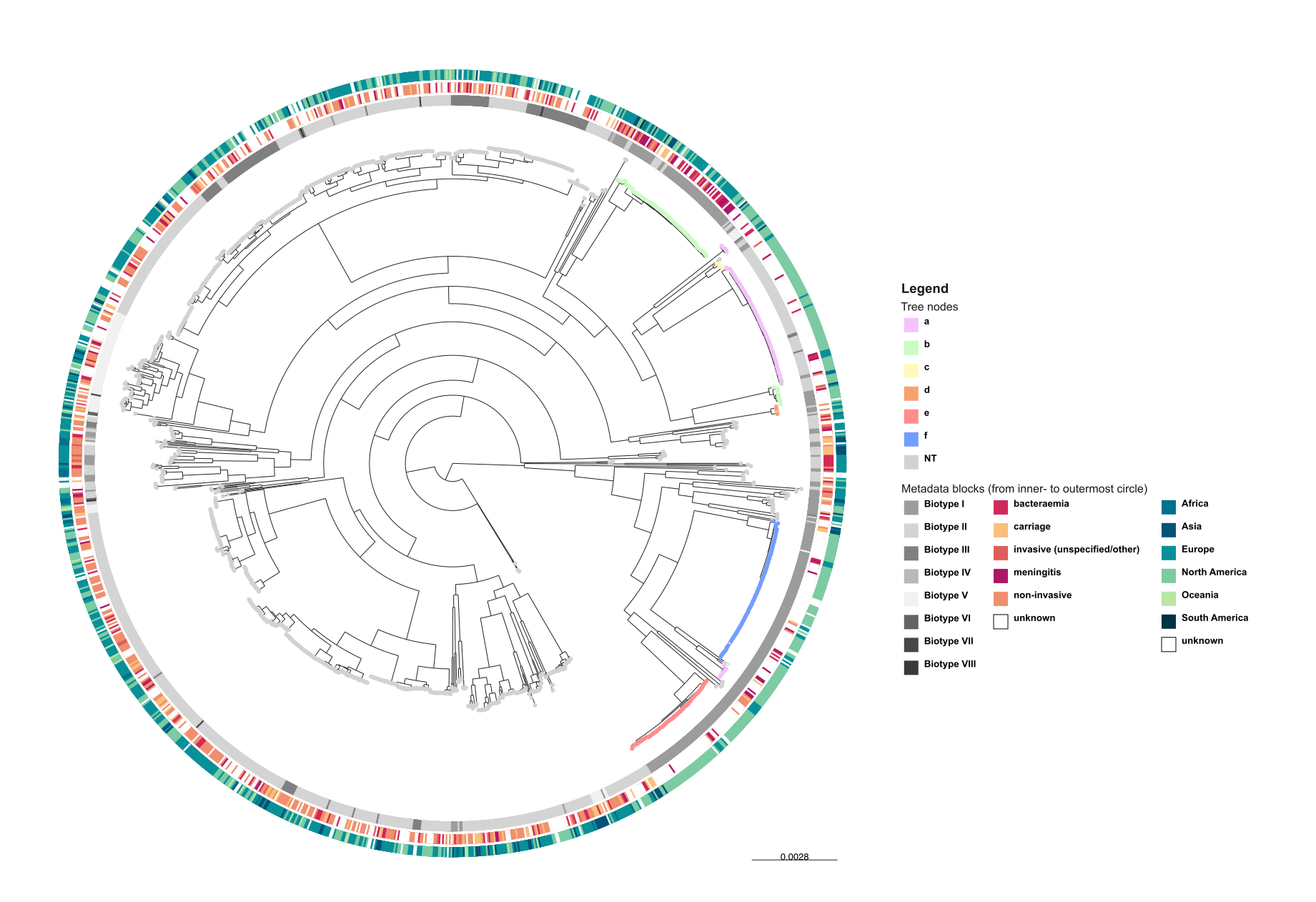
The maximum-likelihood tree from core genome alignment of 1,376 *H. influenzae* genomes in the validation dataset. Tree nodes were coloured based on the capsule type. The innermost metadata block showed biotype assignments, based on the presence/absence of genes encoding ornithine decarboxylase (ODC), urease, and tryptophanase. The middle and outermost metadata block showed disease associated with the isolates and the continent where the isolates originating, respectively.
