## Supplementary Tables for "Development and Implementation of a Core Genome Multilocus Sequence Typing (cgMLST) scheme for *Haemophilus* influenzae"

**Supplementary Table 1. Methods and features of four different pangenome and core genome reconstruction tools.**

| **Tools** | **Description** | **Input files** | **Paralogs detection** | **Pseudogene detection** | **References** |
| --- | --- | --- | --- | --- | --- |
| PIRATE | Rapid investigation of pangenome by classifying orthologous gene families with varied sequence similarity threshold. | Annotation files of draft genomes (.gff) from Prokka | Sequence clustering with CD-HIT and iterative MCL where clusters containing > 1 sequence per genome will go through paralogs classification step, which differentiate fission vs duplicated genes. | None | [1] |
| PEPPAN | Pipeline for pangenome construction which is based on similarity prediction, phylogenetic tree, and synteny. | Fasta and annotation files (.gff, from any source) of draft genomes | Combining tree-based (single BLAST hit, neighbour-joining, or maximum-likelihood algorithm) and synteny-based approach to detect paralogs. | Yes^a^ | [2] |
| chewBBACA | A pipeline with multiple functions to support wg/cgMLST creation and validation which relies on the BSR-based allele calling for validating allele variation. | - Fasta files of draft genomes. - Prodigal training file for bacterial species. | During BSR-based allele calling, each loci is given paralog count (PC) number, which equals to how many times a matching CDS to that loci also matches with other loci. | None | [3] |
| Panaroo | A graph-based clustering tool which corrects annotation error by taking into account information provided by each genome. | Annotation files of draft genomes (.gff) | Sequence clustering at high threshold (98%) with CD-HIT. Clusters with >1 occurrence will be temporarily classified as putative paralogs. The graphical representation of each paralog cluster are assessed using the global context of the graph. | None | [4] |

^a^*Pseudogene is defined as a CDS in genome with a significantly shorter length than other orthologous genes (default setting 0.8, equal or more to this threshold the gene is considered intact).*

^b^*Pseudogene identification can be done by conducting a manual curation.*

*Abbreviation:*

*wgMLST: whole-genome multilocus sequence typing; cgMLST: core genome multilocus sequence typing; MCL: Markov Cluster Algorithm; CDS: coding sequence*

**Supplementary Table 2**

| **Parameters** | **Threshold** | **Rationale for threshold** | **Reference(s)** |
| --- | --- | --- | --- |
| rMLST score | ≥ 85% | A low score for rMLST means uncertainty in species detection. | [5] |
| MLST and rMLST allele designation | One allele per loci | MLST and rMLST loci with multiple alleles suggest impure sequenced isolates and/or poor-quality data. | [5] |
| Genome length | 1.4 Mb – 2.4 Mb | Range of Hinf genome length. A value outside this range may indicate contamination or incorrect species. | [6-8] |
| GC content | 37 – 41mol% | Whole-genome GC content is constant for the same species hence deviation from this range denotes possible contamination. | [9] |
| Number of contigs | ≤ 500 | Lower number of contigs reflect a good-quality sequencing process and genome assembly. | [10] |

**Supplementary Table 3**

| **Clade** | **Gene 1** | **Gene 2** | **Gene 3** | **Gene 4** | **Gene 5** | **Gene 6** | **Gene 7** | **Gene 8** | **Gene 9** | **Gene 10** | **Gene 11** | **Gene 12** | **Gene 13** | **Gene 14** | **Gene 15** | **Gene 16** | **Gene 17** |
| --- | --- | --- | --- | --- | --- | --- | --- | --- | --- | --- | --- | --- | --- | --- | --- | --- | --- |
| NTHi_Clade_I | 1 | 1 | 0 | 2 | 2 | 0 | 0 | 1 | 1 | 1 | 0 | 0 | 0 | 0 | 0 | 0 | 0 |
| NTHi_Clade_II | 1 | 1 | 1 | 0 | 1 | 1 | 1 | 0 | 0 | 0 | 0 | 0 | 0 | 0 | 1 | 1 | 1 |
| NTHi_Clade_II | 1 | 1 | 1 | 0 | 1 | 0 | 1 | 0 | 0 | 0 | 0 | 0 | 0 | 0 | 1 | 0 | 1 |
| NTHi_Clade_II | 1 | 1 | 1 | 0 | 1 | 1 | 1 | 0 | 0 | 0 | 0 | 0 | 0 | 0 | 0 | 1 | 1 |
| NTHi_Clade_III | 1 | 1 | 1 | 1 | 0 | 1 | 1 | 0 | 0 | 0 | 0 | 0 | 0 | 0 | 1 | 1 | 1 |
| NTHi_Clade_III | 1 | 1 | 1 | 1 | 0 | 0 | 1 | 0 | 0 | 0 | 0 | 0 | 0 | 0 | 1 | 0 | 1 |
| NTHi_Clade_III | 1 | 1 | 1 | 1 | 0 | 1 | 1 | 0 | 0 | 0 | 0 | 0 | 0 | 0 | 0 | 1 | 1 |
| NTHi_Clade_IV | 0 | 0 | 1 | 2 | 2 | 1 | 1 | 0 | 0 | 0 | 1 | 1 | 1 | 1 | 1 | 1 | 0 |
| NTHi_Clade_IV | 0 | 0 | 1 | 2 | 2 | 0 | 1 | 0 | 0 | 0 | 1 | 1 | 1 | 1 | 1 | 0 | 0 |
| NTHi_Clade_IV | 0 | 0 | 1 | 2 | 2 | 1 | 1 | 0 | 0 | 0 | 1 | 1 | 1 | 1 | 0 | 1 | 0 |
| NTHi_Clade_V | 1 | 1 | 1 | 2 | 1 | 1 | 1 | 0 | 0 | 0 | 0 | 0 | 0 | 0 | 1 | 1 | 0 |
| NTHi_Clade_V | 1 | 1 | 1 | 2 | 1 | 0 | 1 | 0 | 0 | 0 | 0 | 0 | 0 | 0 | 1 | 0 | 0 |
| NTHi_Clade_V | 1 | 1 | 1 | 2 | 1 | 1 | 1 | 0 | 0 | 0 | 0 | 0 | 0 | 0 | 0 | 1 | 0 |
| NTHi_Clade_VI | 1 | 1 | 1 | 2 | 2 | 0 | 1 | 0 | 0 | 0 | 0 | 0 | 0 | 0 | 0 | 0 | 2 |

**Note:**

0: absence

1: presence

2: presence or absence

| **Gene** | **Common names[11]** | **PubMLST loci name** |
| --- | --- | --- |
| Gene 1 | HIFGL_RS07705 | HAEM0066 |
| Gene 2 | HIFGL_RS07710 | HAEM0067 |
| Gene 3 | HIB_RS06975 | HAEM1319 |
| Gene 4 | HIB_RS07380 | HAEM1400 |
| Gene 5 | HIFGL_RS06855 | HAEM1857 |
| Gene 6 | HMWC | HAEM1904 |
| Gene 7 | HIFGL_RS03555 | HAEM1974 |
| Gene 8 | HIFGL_RS05025 | HAEM1978 |
| Gene 9 | HIFGL_RS05250 | HAEM1979 |
| Gene 10 | HIFGL_RS07070 | HAEM1980 |

| **Gene** | **Common names[11]** | **PubMLST loci name** |
| --- | --- | --- |
| Gene 11 | C645_RS00655 | HAEM1981 |
| Gene 12 | C645_RS00650 | HAEM1982 |
| Gene 13 | C645_RS00645 | HAEM1983 |
| Gene 14 | C645_RS08170 | HAEM1984 |
| Gene 15 | HMWA | HAEM1985 |
| Gene 16 | HMWB | HAEM1986 |
| Gene 17 | R2846_RS08405 | HAEM1987 |

**Supplementary Table 4.1**

| **Biotypes** | **Indole** | **Urease** | **ODC** |
| --- | --- | --- | --- |
| Biotype I | 1 | 1 | 1 |
| Biotype II | 1 | 1 | 0 |
| Biotype III | 0 | 1 | 0 |
| Biotype IV | 0 | 1 | 1 |
| Biotype V | 1 | 0 | 1 |
| Biotype VI | 0 | 0 | 1 |
| Biotype VII | 1 | 0 | 0 |
| Biotype VIII | 0 | 0 | 0 |

**Note:**

ODC: ornithine decarboxylase

0: absence

1: presence

**Supplementary Table 4.2**

| **Enzymes/products** | **NCBI gene names[12]** | **PubMLST loci name** | **Note on NCBI gene names** |
| --- | --- | --- | --- |
| Indole | HI_1389.1 | HAEM1528 | Original gene record was discontinued.  New gene record: https://www.ncbi.nlm.nih.gov/gene/72525313 (from NCBI reference genome 477) |
| Urease | HI_053 | HAEM0653 | Original gene record was suppressed.  New gene record: https://www.ncbi.nlm.nih.gov/gene/72526651 (from NCBI reference genome 477) |
| ODC | HI_0590 | HAEM0709 | Original gene record was suppressed.  New gene record: https://www.ncbi.nlm.nih.gov/gene/12596673 (from NCBI complete genome R2866) |

**Supplementary Table 5.1**

| **PubMLST ID** | **Isolate ID** | **BioSample accession** | **Capsule genotype** | **Year of collection** | **Clinical source; location [Reference PMID]** |
| --- | --- | --- | --- | --- | --- |
| 1 | 477 | SAMN02595602 | NTHi | 1994 | middle ear fluid, otitis media; Finland [12682154] |
| 495 | R3159/2019 | SAMN03450976 | NTHi | N/A | sputum, chronic obstructive pulmonary disease; USA [1581306] |
| 2222 | 10P129H1 | SAMN09258515 | NTHi | 2005 | sputum, chronic obstructive pulmonary disease; USA [30533802] |
| 5068 | 86-028NP | SAMN02603157 | NTHi | 1987 | NP swab, other; USA [15968074;22377449;24706866;28604066] |
| 5069 | R2846 | SAMN02604260 | NTHi | 2010 | middle ear fluid, otitis media; USA [22377449] |
| 5082 | 10810 | SAMEA3138383 | b | 2010 | CSF, meningitis; USA [22377449] |
| 5083 | F3031 | SAMEA3138342 | NTHi | 1984 | other, Brazilian puerpuric fever; Brazil [22377449;24706866] |
| 5084 | F3047 | SAMEA2272570 | NTHi | 1984 | eye, conjunctivitis associated with Brazilian puerpuric fever; Brazil [22377449;24706866] |
| 5230 | Hi375 | SAMEA1411782 | NTHi | 1995 | middle ear fluid, otitis media; Finland [24706866] |
| 5257 | R2866 | SAMN02604259 | NTHi | 1996 | CSF, meningitis; USA [22377449;24706866;28604066] |
| 5284 | CGSHiCZ412602 | SAMN02800345 | NTHi | 2014 | middle ear fluid, otitis media; Czech Republic [23865594] |
| 5559 | 723 | SAMN02595605 | NTHi | 1994 | middle ear fluid, otitis media; Finland [28604066] |
| 5571 | C486 | SAMN02595604 | NTHi | 2014 | middle ear fluid, otitis media; USA [28604066] |
| 19836 | 84P36H1 | SAMN09258517 | NTHi | 2000 | sputum, chronic obstructive pulmonary disease; USA [30533803] |

**Supplementary Table 5.2**

| **PubMLST ID** | **Contigs** | **Total length** | **N50** | **L50** | **No of loci annotated in PubMLST** | **Sequencing technology** |
| --- | --- | --- | --- | --- | --- | --- |
| 1 | 1 | 1,846,259 | 1,846,259 | 1 | 1591 | PacBio |
| 495 | 1 | 1,969,659 | 1,969,659 | 1 | 1571 | Sanger dideoxy sequencing; 454 |
| 2222 | 1 | 2,047,595 | 2,047,595 | 1 | 1152 | PacBio |
| 5068 | 1 | 1,914,490 | 1,914,490 | 1 | 1633 | n/a |
| 5069 | 1 | 1,819,370 | 1,819,370 | 1 | 1550 | n/a |
| 5082 | 1 | 1,981,535 | 1,981,535 | 1 | 1907 | n/a |
| 5083 | 1 | 1,985,832 | 1,985,832 | 1 | 1525 | n/a |
| 5084 | 1 | 2,007,018 | 2,007,018 | 1 | 1456 | n/a |
| 5230 | 37 | 1,815,459 | 145,197 | 5 | 1568 | PacBio |
| 5257 | 1 | 1,932,306 | 1,932,306 | 1 | 1621 | n/a |
| 5284 | 1 | 1,811,802 | 1,811,802 | 1 | 1572 | PacBio RSII |
| 5559 | 46 | 1,851,915 | 94,087 | 7 | 1564 | PacBio |
| 5571 | 1 | 1,846,503 | 1,846,503 | 1 | 1574 | PacBio |
| 19836 | 1 | 2,025,527 | 2,025,527 | 1 | 1277 | Illumina HiSeq 2000 |

**Supplementary Table 6**

| Capsule type | CGC group at 500 allelic mismatch | Clonal complex(es) |
| --- | --- | --- |
| a | 46 | NA |
|  | **48*** | ST-23 complex |
|  | 79 | NA |
|  | 81 | NA |
| b | 3* | ST-6 complex |
|  | 78 | ST-222 complex |
| c | 58 | ST-7 complex |
| d | **70** | ST-10 complex |
| e | 40* | ST-18 complex |
| f | **38*** | **ST-124 complex#** |
|  |  | ST-123 complex |
|  |  | **ST-210 complex** |
|  |  | **ST-422 complex** |
|  | 129 | ST-123 complex |
| NT | 2 | ST-321 complex |
|  | 6 | ST-393 complex |
|  | 9 | ST-84 complex |
|  | 11 | ST-472 complex |
|  | 12 | ST-584 complex |
|  | 18 | ST-2118 complex |
|  | 20 | ST-393 complex |
|  | 28 | ST-11 complex |
|  | 29 | ST-836 complex |
|  | 30 | **ST-422 complex** |
|  | 32 | ST-139 complex |
|  | 34 | ST-264 complex |
|  | 35 | ST-165 complex |
|  | 36 | ST-425 complex |
|  | 37 | ST-3 complex |
|  | **38** | **ST-124 complex** |
|  | 41 | ST-1426 complex |
|  | 42 | ST-266 complex |
|  | **48** | ST-23 complex |
|  | 59 | ST-986 complex |
|  | 61 | ST-249 complex |
|  | 67 | ST-746 complex |
| **Supplementary Table 5 cont’** | | |
| Capsule type | CGC group at 500 allelic mismatch | Clonal complex(es) |
| NT | **70** | ST-10 complex |
|  | 73 | **ST-210 complex** |
|  | 84 | ST-385 complex |
|  | 88 | ST-262 complex |
|  | 95 | ST-1736 complex |
|  | 97 | ST-1030 complex |
|  | 106 | ST-1477 complex |
|  | 107 | ST-589 complex |
|  | 108 | ST-84 complex |
|  | 110 | ST-474 complex |
|  | 125 | **ST-210 complex** |
|  | 127 | ST-706 complex |
|  | 137 | **ST-210 complex** |
|  | 143 | ST-1030 complex |
|  | 159 | **ST-210 complex** |
|  | 1 | ST-3 complex |
|  |  | ST-389 complex |
|  | 10 | ST-34 complex |
|  |  | ST-396 complex |
|  | 14 | ST-1836 complex |
|  |  | ST-41 complex |
|  | 16* | ST-1477 complex |
|  | 22 | ST-105 complex |
|  |  | ST-163 complex |
|  | 25 | **ST-422 complex** |
|  |  | ST-57 complex |
|  | 43* | ST-513 complex |
|  | 56* | ST-519 complex |
|  | 117* | ST-270 complex |
|  | 139* | ST-3 complex |
|  | 5* | ST-139 complex |
|  |  | ST-3 complex |
|  | 7 | ST-1025 complex |
|  |  | ST-199 complex |
|  |  | ST-652 complex |
|  | 26 | ST-163 complex |
| **Supplementary Table 5 cont’** | | |
| Capsule type | CGC group at 500 allelic mismatch | Clonal complex(es) |
| NT |  | ST-3 complex |
|  |  | ST-487 complex |
|  | 27* | ST-1529 complex |
|  |  | ST-155 complex |
|  | 23* | ST-107 complex |
|  |  | ST-390 complex |
|  |  | ST-395 complex |
|  | 24* | ST-139 complex |
|  |  | ST-142 complex |
|  |  | ST-3 complex |
|  | 15* | ST-12 complex |
|  |  | **ST-124 complex** |
|  |  | ST-183 complex |
|  |  | ST-3 complex |
|  | 31* | ST-210 complex |
|  |  | ST-242 complex |
|  |  | ST-430 complex |
|  |  | ST-931 complex |
|  | 44 | NA |
|  | 50 | NA |
|  | 51 | NA |
|  | 54 | NA |
|  | 55 | NA |
|  | 62 | NA |
|  | 64 | NA |
|  | 65 | NA |
|  | 71 | NA |
|  | 74 | NA |
|  | 75 | NA |
|  | 77 | NA |
|  | 82 | NA |
|  | 85 | NA |
|  | 86 | NA |
|  | 87 | NA |
|  | 89 | NA |
|  | 90 | NA |
| **Supplementary Table 5 cont’** | | |
| Capsule type | CGC group at 500 allelic mismatch | Clonal complex(es) |
| NT | 91 | NA |
|  | 92 | NA |
|  | 94 | NA |
|  | 99 | NA |
|  | 100 | NA |
|  | 101 | NA |
|  | 102 | NA |
|  | 104 | NA |
|  | 105 | NA |
|  | 109 | NA |
|  | 111 | NA |
|  | 112 | NA |
|  | 113 | NA |
|  | 115 | NA |
|  | 118 | NA |
|  | 119 | NA |
|  | 120 | NA |
|  | 121 | NA |
|  | 122 | NA |
|  | 123 | NA |
|  | 124 | NA |
|  | 126 | NA |
|  | 128 | NA |
|  | 142 | NA |
|  | 144 | NA |
|  | 148 | NA |
|  | 149 | NA |
|  | 150 | NA |
|  | 153 | NA |
|  | 156 | NA |
|  | 157 | NA |
|  | 165 | NA |

| * | There are genomes within this CGC group without clonal complex assigned. |
| --- | --- |
| **Black bold text** | This CGC group is assigned to both typeable and non-typeable *H. influenzae*. The NTHi genome with this CGC group is clustered with the typeable *H. influenzae* genomes, a possible observation of rare capsule loss events. [ref] |
| # | The ST-124 complex is the major clonal complex associated with CGC500 38 (92/97 genomes within the GCG group). |
| Red text, not bolded | NTHi genomes with this clonal complex can belong to different CGC500 groups. This underlines that clustering based on the 7-loci MLST does not reflect a phylogenetic relatedness, especially for NTHi. |
| **Bold red text** | This clonal complex is assigned to both typeable and non-typeable. [may need to add explanation later] |
